## Supplementary material for "The neuropeptide sulfakinin is a peripheral regulator of insect behavioral switch between mating and foraging": Chemicals used for electroantennograms

**Supplementary File 1. Chemicals used for electroantennograms.**

| **Source** | **Order** | **Chemicals** | **Vendor** | **Purity** | **CAS#** |
| --- | --- | --- | --- | --- | --- |
| Food odors | 1 | Ethyl benzoate | Sigma-Aldrich | 99% | 93-89-0 |
|  | 2 | Diethyl maleate | Sigma-Aldrich | 97% | 141-05-9 |
|  | 3 | Ethyl butyrate | Sigma-Aldrich | 99% | 105-54-4 |
|  | 4 | Geraniol | Sigma-Aldrich | 98% | 106-24-1 |
|  | 5 | 1-Octen-3-ol | Sigma-Aldrich | 98% | 3391-86-4 |
|  | 6 | 4-Carvomenthenol | Sigma-Aldrich | 95% | 562-74-3 |
|  | 7 | Citronellal | Sigma-Aldrich | 95% | 106-23-0 |
|  | 8 | Trans-2-Hexenal | Aladdin | 98% | 6728-26-3 |
|  | 9 | ME | Sigma-Aldrich | 99% | 93-15-2 |
|  | 10 | β-Caryophyllene | Aladdin | 98% | 87-44-5 |
|  | 11 | (R)-(+)-Limonene | Sigma-Aldrich | 97% | 5989-27-5 |
|  | 12 | Dimethylacetic acid | Sigma-Aldrich | 99% | 79-31-2 |
| Sex pheromones | 13 | 2,3,5-trimethylpyrazine | Aladdin | 98% | 14667-55-1 |
|  | 14 | 2,3,5,6-tetramethylpyrazine | Aladdin | 98% | 1124-11-4 |
|  | 15 | Ethyl laurate | Aladdin | 99% | 106-33-2 |
|  | 16 | Ethyl myristate | Aladdin | 98% | 124-06-1 |
|  | 17 | Ethyl palmitate | Aladdin | 98% | 628-97-7 |
