## Supplementary material for "The neuropeptide sulfakinin is a peripheral regulator of insect behavioral switch between mating and foraging": Primer sequences used in this study

**Supplementary File 2. Primer sequences used in this study**

| **Experiments** | **Gene name** | **Primer name** | **Primer sequences (5′ → 3′)** |
| --- | --- | --- | --- |
| RT-qPCR | RPS3 | q-RPS3-F | TAAGTTGACCGGAGGTTTGG |
|  |  | q-RPS3-R | TGGATCACCAGAGTGGATCA |
|  | α-tubulin | q-α-tubulin-F | CGCATTCATGGTTGATAACG |
|  |  | q-α-tubulin-R | GGGCACCAAGTTAGTCTGGA |
|  | CCHamide1 | q-CCHamide1-F | CAAGCTTGTTACGGTGGACA |
|  |  | q-CCHamide1-R | TCAACAGGTCGCTTCAGATG |
|  | CCHamide1R | q-CCHamide1-F | GCTTAAGCCACGACTCGAAC |
|  |  | q-CCHamide1-R | ATATCCACCACCCTCATCCA |
|  | GPB5 | q-GPB5-F | ATCCCGTTTGTGTACATGCG |
|  |  | q-GPB5-R | ATATTGTTCACCGGCGCTTC |
|  | GPB5R | q-GPB5R-F | CGCTGCAAAAAGAATGACAA |
|  |  | q-GPB5R-R | AAATCGGTTCGGGCTAAACT |
|  | ITG-like peptide | q- ITG-like peptide -F | CAATACGGACTGTGAGCTGG |
|  |  | q- ITG-like peptide -R | CGACGAGACTGCCACCATC |
|  | ITP1 | q-ITP1-F | GCCACACAACAACCACAATC |
|  |  | q-ITP1-R | GGCATTCTCCAAACCATTTC |
|  | ITP2 | q-ITP2-F | GCCACACAACAACCACAATC |
|  |  | q-ITP2-R | TCCATCTCTTCGTGCAGTTG |
|  | Lecokinin | q-Lecokinin-F | GGACCAACAAGACTCCGGTA |
|  |  | q-Lecokinin-R | GCTTCTTACCCAATACAACGG |
|  | LecokininR1 | q-LecokininR1-F | CGAGTCGGAATGGCAGTAAT |
|  |  | q-LecokininR1-R | TAGGTGTCGAGTGCATCAGC |
|  | LecokininR2 | q-LecokininR2-F | CACTGGCATCGTGGTGTTAC |
|  |  | q-LecokininR2-R | AAAGCGGCTTGAAACTGAAA |
|  | SIFamide | q-SIFamide-F | CCGTATGTTTCGTAGCCTTGA |
|  |  | q-SIFamide-R | AATTTCGCACATTGCAGTCA |
| **Experiments** | **Gene name** | **Primer name** | **Primer sequences (5′ → 3′)** |
| RT-qPCR | SIFamideR1 | q-SIFamideR1-F | TGCAACAGAAGTCGAAGGTG |
|  |  | q-SIFamideR1-R | CGAAGCCACGTCGATATTTT |
|  | SIFamideR2 | q-SIFamideR2-F | TATGCGCAACTCCACCAATA |
|  |  | q-SIFamideR2-R | TGAGATGGCCAAAATTGTCA |
|  | SIFamideR3 | q-SIFamideR3-F | TGTGCCAGCGCTAGATATTG |
|  |  | q-SIFamideR3-R | GGAGTACGGTGTCCAGGCTA |
|  | Sulfakinin | q-Sulfakinin-F | TGGTGGCCTTAACGTTGACT |
|  |  | q-Sulfakinin-R | CCAGAGGCATACCACCAGAT |
|  | SulfakininR1 | q-SulfakininR1-F | TCAAACGAGGCGAAAAATCT |
|  |  | q-SulfakininR1-R | CGTAAATAGCTGGGCCGATA |
|  | SulfakininR2 | q-SulfakininR2-F | TATTGTTGGGGGTCTTCTGC |
|  |  | q-SulfakininR2-R | ATAGCGTTCGCAGGATATGG |
|  | NPF1 | q-NPF1-F | CTGCCGCTGACGAACCTTTTT |
|  |  | q-NPF1-R | TAATCCCCACGGAATCCTTTGC |
|  | NPFR | q-NPFR-F | TTTTCGTCTCGACCATTTCC |
|  |  | q-NPFR-R | TTCGACGCAGAAGCATACAC |
|  | Bur-beta-subunit | q-Bursicon -beta-subunit-F | TCTTTGTGCTGAGTCGGTTG |
|  |  | q-Bursicon -beta-subunit-R | ACGGACGGCTGTACTTGACT |
|  | Bur-beta-subunitR | q-Bursicon -beta-subunitR-F | CAAACTCAGGCGAGGAGAAC |
|  |  | q-Bursicon -beta-subunitR-R | GCTTATTTGTGCCACCACCT |
|  | NPLP | q-NPLP-F | CGTGCCGGACTATGACTACA |
|  |  | q-NPLP-R | GCCCACATAACGCTTGTCTT |
|  | Tachykinin | q-Tachykinin-F | CAATCTGGCAATGGATGATG |
|  |  | q-Tachykinin-R | ACGCAAACGGTCCTCTTCTA |
| **Experiments** | **Gene name** | **Primer name** | **Primer sequences (5′ → 3′)** |
| RT-qPCR | TachykininR | q-TachykininR-F | ACCCGAATTGTATGCTACGC |
|  |  | q-TachykininR-R | TGCCGACTCGTGAATAAGTG |
|  | sNPF | q-sNPF-F | ATGTTGCTCTCACTCCGTCA |
|  |  | q-sNPF-R | GTTGTTGTTGTGTTGCTGGC |
|  | sNPFR | q-sNPFR-F | GTGGCTTCGGTAACTGGTGT |
|  |  | q-sNPFR-R | TGTGCGCTTCTTACGTTCAC |
|  | Natalisin | q-Natalisin-F | GACAGAGCGAATGAGGTGAA |
|  |  | q-Natalisin-R | CAGCAGACGACTGATAGCAAC |
|  | NatalisinR | q-NatalisinR-F | TGTGGATTGTTGCTGGTCAT |
|  |  | q-NatalisinR-R | AATAGAACCGAAGGGCCAAT |
|  | OR7a.4 | q-OR7a.4-F | GCAGCACCGCATATCTTCAA |
|  |  | q-OR7a.4-R | CACCCAAATGACTAACGCGT |
|  | OR7a.8 | q-OR7a.8-F | TGAACAGGATCGGCGTTTTC |
|  |  | q-OR7a.8-R | TTCCTGACCAATGCCCATCT |
|  | OR10a | q-OR10a-F | GTACTTTTCCACTGCGCGAT |
|  |  | q-OR10a-R | GTGTCCAAAGCGAGTTCCAG |
|  | OR49a | q-OR49a-F | GCATGGATAACTCAAAGGCAGA |
|  |  | q-OR49a-R | CGAAACCAAAGCCCTCCAAT |
|  | OR63a | q-OR63a.2-F | GGACATGTTTTTCGGTGCTT |
|  |  | q-OR63a.2-R | CTGGACCAGTCCACAGGAAT |
|  | OR67c.1 | q-OR67c.1-F | TTTAAGAGCAGTGAACCCGC |
|  |  | q-OR67c.1-R | AGATGTCGCTAACTCGTCGT |
|  | OR67d.3 | q-OR67d.3-F | ATCTTGTGAACGTGTGTGGC |
|  |  | q-OR67d.3-R | GTGTCGACATCAATGCCAGG |
|  | OR74a | q-OR74a-F | TCGCCAAATCAATGCAGGAG |
|  |  | q-OR74a-R | GGCGTCCAAAGATCAGCAAT |
|  | OR94b.1 | q-OR94b.1-F | AAGTGGCCAACGATCCCATA |
|  |  | q-OR94b.1-R | CATCGGCTGGTGTCATCATG |
| **Experiments** | **Gene name** | **Primer name** | **Primer sequences (5′ → 3′)** |
| RT-qPCR | Sk | SK-F | TTGAGCATTCAAAACGAAGC |
|  |  | SK-R | AAACATTTTCGGGCCGTAG |
| Off-target analysis | Sk-potential off-target-1 | SK-OFF1-F | CTAAATCTCCGCGAAGCAAC |
|  |  | SK-OFF1-R | TTTTGAAAGCCGGATAATGC |
|  | Sk-potential off-target-2 | SK-OFF2-F | GCGTCTAAGTGCCAATGTGA |
|  |  | SK-OFF2-F | CAAGGAGCTGGTGTTGTTTG |
|  | SkR1 | SKR1-F | GCGAACTGTTGAACGAGGAT |
|  |  | SKR1-R | AGCAGTGTGCCGACTAAGGT |
|  | SkR1-potential off-target-1 | SKR1-OFF1-F | CCTCTGCGTCAACTCGTACA |
|  |  | SKR1-OFF1-R | GCTAGCTGGGAGGCTATCAA |
|  | SkR1-potential off-target-2 | SKR1-OFF2-F | GAACAAAAGCCAATGCAACA |
|  |  | SKR1-OFF2-R | CCACTAATCATTAAACGGCG |
|  | SkR1-potential off-target-3 | SKR1-OFF3-F | TGAACACTTTGGCAGTTTGG |
|  |  | SKR1-OFF3-R | TTGACGCGTGTAAAATCTGC |
