## Supplementary material for "The neuropeptide sulfakinin is a peripheral regulator of insect behavioral switch between mating and foraging": Analysis of off-target effects of mutants

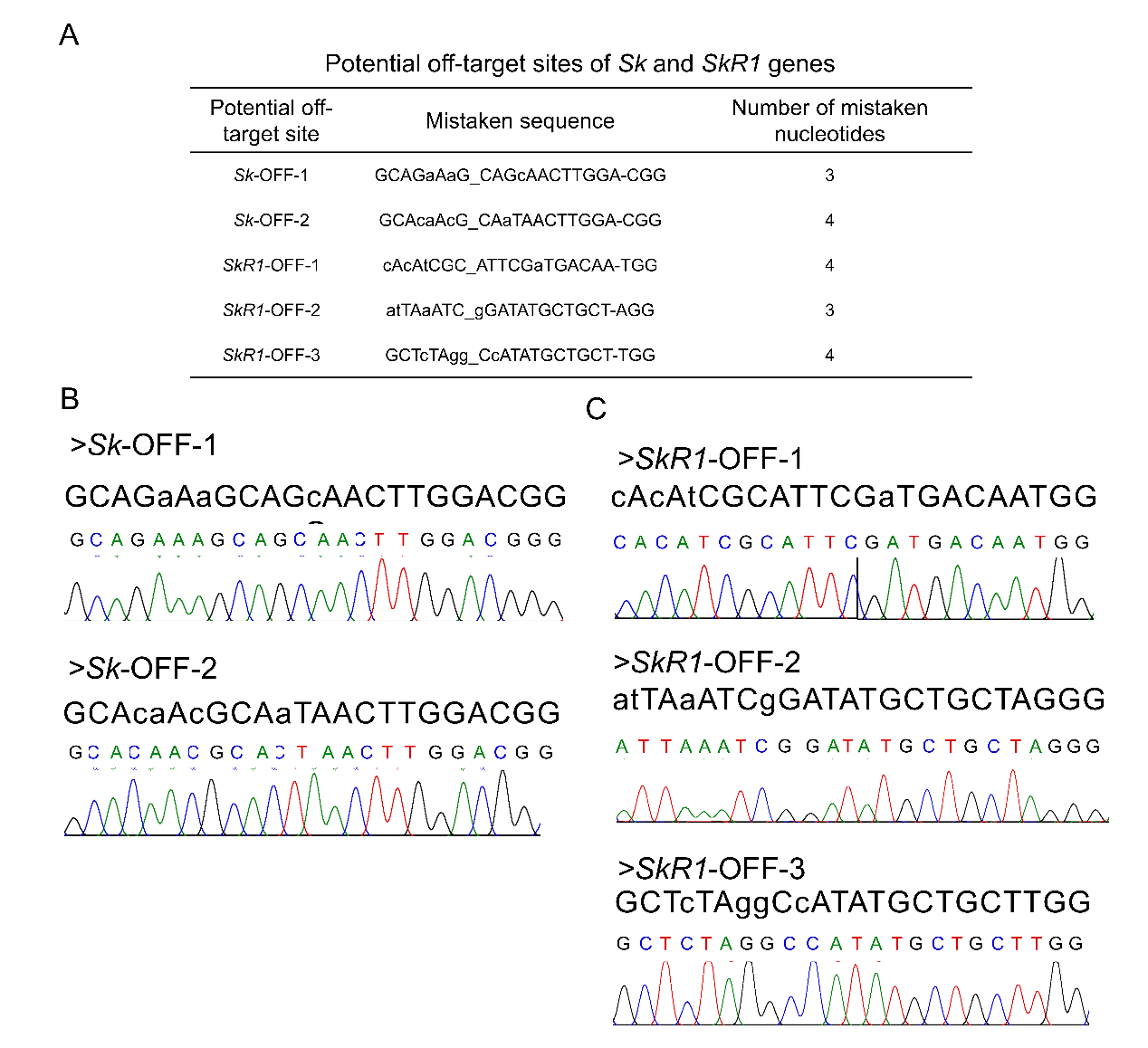

**Supplementary File 3.** Analysis of off-target effects of mutants. (*A*) Prediction of potential off-target sites of the *Sk* and *SkR1* genes. (*B*) PCR amplification and sequencing of potential off-target sites of the *Sk* gene. (*C*) PCR amplification and sequencing of potential off-target sites of the *SkR1* gene.
